# “Lactic acid influences iron assimilation by a fungal pathogen via the iron reductive uptake pathway”

**DOI:** 10.1101/2025.05.27.656276

**Authors:** Alexandra Gomes-Gonçalves, Wouter Van Genechten, Patrícia Ataíde, Cláudia Barata-Antunes, Faezeh Ghasemi, Margarida Casal, Miguel C. Teixeira, Joaquín Ariño, Alistair J.P. Brown, Patrick Van Dijck, Sandra Paiva

## Abstract

*Candida albicans* is a fungal commensal of humans that often causes mucosal infections in otherwise healthy individuals, and also serious infections in immunocompromised patients. The capacity of this fungus to colonise and cause disease relies on its ability to grow within the host, adapting to various nutrient restrictions and physicochemical conditions. The presence of alternative carbon sources, such as the lactate produced by the local microbiota, influences *C. albicans* antifungal drug resistance and immune evasion. In this study, we used genome-wide transcriptomic analysis to investigate the effect of lactate exposure upon metabolic rewiring. We provide evidence that *C. albicans* cells respond to growth in the presence of lactate at pH 5 by regulating genes encoding micronutrient transporters, notably iron transporters. More specifically, lactate triggers the downregulation of genes on the reductive iron uptake pathway, inferring a diminished requirement for high-affinity iron uptake. This is supported by the observation that lactate promotes the intracellular accumulation of iron by *C. albicans* cells. Lactate even enhances the growth of iron-transport defective *C. albicans* cells under iron-limited conditions. Lactate is known to activate protein kinase A (PKA) signalling. However, lactate-induced iron assimilation is PKA-independent. This work provides new insights into the role of lactate in iron homeostasis – two important factors that promote *C. albicans* virulence in the mammalian host, where nutritional immunity is a key antimicrobial strategy.

**Importance:** *Candida albicans* is a major opportunistic fungal pathogen capable of causing life- threatening infections, particularly in immunocompromised individuals. Its ability to adapt to diverse host environments underlies its success as a commensal and pathogen. This study provides new insights into the metabolic flexibility of *C. albicans*, with a specific focus on how lactate, a common carbon source in host niches, influences iron acquisition and homeostasis. Our findings reveal that, during growth at pH 5, lactate modulates the expression of micronutrient transporters and enhances iron assimilation in *C. albicans*. These results suggest a role of lactate in promoting iron uptake, potentially facilitating fungal colonization and persistence within the host. By elucidating the molecular and phenotypic consequences of lactate exposure upon iron metabolism, this study contributes to a deeper understanding of host-pathogen interactions.

## Introduction

*Candida albicans,* a major fungal pathogen, normally resides as a commensal organism in the microbiota of most humans (1, 2). However, when the microbiota is disrupted, *C. albicans* can cause mucosal infections and, in immunocompromised patients, potentially fatal systemic infections with mortality rates of up to 50 % (3, 4). The ability of *C. albicans* to colonise and infect its host relies on its capacity to rapidly adjust to diverse niches in the human body, each presenting distinct nutritional conditions (5). A key aspect of this adaptability is its metabolic flexibility, particularly its ability to use alternative carbon sources, such as the carboxylic acids lactate and acetate. The abundances of these carboxylic acids vary between host environments like the gastrointestinal tract, vaginal mucosa, and bloodstream (6–9). *C. albicans* exploits specific transport proteins to assimilate carboxylic acids, such as the Jen1 and Jen2 carboxylate/proton symporters (10, 11). Ato proteins, which are members of the Acetate Uptake Transporter (AceTr) family, have also been implicated in acetate transport and contribute to metabolic adaptability (12). This enables *C. albicans* to thrive under nutrient-limited conditions and contributes to the virulence of this fungus both directly and indirectly.

Numerous studies have examined the influence of carboxylic acids on *C. albicans*. For example, lactate impacts biofilm formation, sensitivity to stresses and antifungal drug resistance in *C. albicans* (6–9). Additionally, exposure to lactate induces changes in cell wall architecture, thereby enhancing *C. albicans* virulence and immune evasion (9). More specifically, lactate sensing enhances the removal of exposed β-glucans, partly through up-regulation of secreted glucanases Xog1 and Eng1 in a protein kinase A (PKA) dependent manner (13–15).

Lactic acid has also been reported to shift the oxidation state of iron from Fe^2+^ to Fe^3+^ (16). This shift in oxidation state may affect the assimilation by *C. albicans* of iron, an essential micronutrient. In *C. albicans,* the iron reductive uptake pathway involves two sequential modifications of the iron oxidation state. Firstly, ferric (Fe^3+^) ions are reduced to ferrous (Fe^2+^) ions by ferric reductases to liberate iron from host sequestration mechanisms (17). Subsequently, Fe^2+^ is re-oxidized to Fe^3+^ by multicopper oxidases. This oxidation step is coupled to the iron permeases, Ftr1 and Ftr2, thereby resulting in the internalisation of the iron (18, 19). Therefore, by influencing iron oxidation state, the presence of lactic acid could, in principle, counteract the initial reduction of Fe^3+^ to Fe^2+^ by ferric reductases, and/or bypass the need for subsequent Fe^2+^ re-oxidation by multicopper oxidases. Indeed, exposure of *C. albicans* to high concentrations of lactic acid has been reported to adversely affect iron uptake and thus induce the expression of genes involved in the reductive iron pathway (20). Interestingly, iron deprivation also induces β-glucan masking (21), a process that, as described earlier, has also been attributed to *C. albicans* exposure to lactate. Both lactate and iron deprivation-induced β-glucan masking are dependent on PKA signalling (21). Taken together, these observations thereby suggest a potential interplay between lactic acid and iron signalling converging on the PKA pathway.

We examine this potential interplay in this study. We show that growth in the presence of physiologically relevant concentrations of lactic acid at pH 5 results in the downregulation of *C. albicans* genes involved in the reductive iron uptake pathway, and the concomitant intracellular accumulation of iron. This intracellular accumulation is dependent upon the iron reductive uptake pathway, but independent of PKA signalling. Our data suggest that, as well as providing a carbon source, the presence of lactate in host niches enhances fungal colonisation by enhancing iron assimilation.

## Results and Discussion

### pH-dependent transcriptional regulation of metal ion transport in response to lactate

*C. albicans* cells are frequently grown under laboratory conditions that do not closely mimic physiological environments, leaving potential physiological effects largely unexplored. To understand the impact of lactate-enriched environments on *C. albicans*, we performed genome-wide transcriptional profiling of *C. albicans* planktonic cells by RNA sequencing. Cells were grown in RPMI media containing low glucose (0.2 %), either at pH 5 or 7, from an initial OD600nm of 0.3. When the cultures reached an OD600nm of 1.0, they were supplemented with 0 % or 0.5 % lactic acid (t = 0; Figure 1A). Cells were then harvested for RNA sequencing at various timepoints up to 4 h. A total of 30 independent samples were collected for the four conditions: n=3 at 1 h; and n = 1 at 0.5, 2, and 4 h. This sampling strategy permitted robust comparison of the effects of lactate and pH upon gene expression at the 1 h timepoint, while revealing trends in gene expression over the 4 h time course.

**Figure 1.**
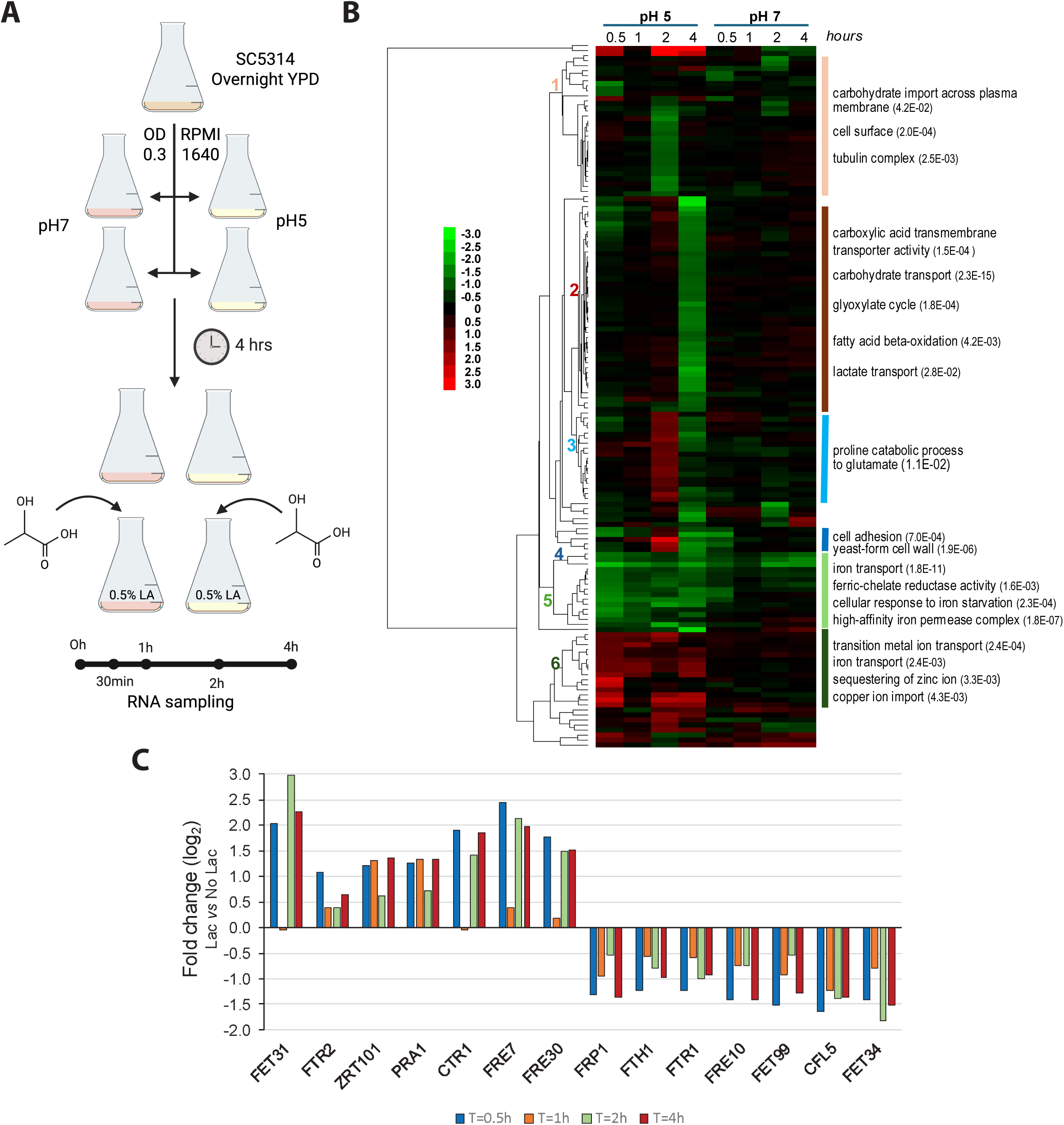
Time profile of genes induced in the presence of lactate, at pH 5 and 7. **(A)** Schematic representation of the experimental setup. Cells were grown in RPMI pH 5.0 or pH 7.0; after 4h and reaching an OD of 1, lactate was added to the cell cultures at a final concentration of 0.5 % (v/v). Samples were collected at 0h, 30 min, 1h, 2h, and 4h after lactate addition. **(B)** Heat map of genes differently expressed in the presence of lactate, either at pH 5 or pH 7. The numbers on the left identify the main clusters, with relevant genes represented on the right. Numbers in parentheses denote the p-value for each specific GO term enrichment. **(C)** Fold change (log2) of genes differently expressed in the presence of lactate related to iron, zinc, and copper transport (pH 5.0) based on the transcriptome analysis

The RNA sequencing yielded at least 10 million aligned reads per sample (mean ± SD, 16.1 ± 4.6 million). Pair-wise comparisons were performed between cells cultured in the presence or absence of lactate for each time point and both pHs. One hundred forty genes were differentially affected by the presence of lactate at least at one time- point in cells grown at pH 5 or 7 (Figure S1). Among them, 131 were differentially expressed at pH 5 (45 were induced, whereas 91 were repressed). The influence of lactate was much less prominent at pH 7: only 21 DEGs, of which 16 were repressed and five induced by lactate. As shown in Figure S1, only one gene was induced by lactate both at pH 5 and 7, whereas about 50 % of genes repressed at pH 7 were also repressed at pH 5.

Transcriptional changes induced by lactate were subjected to hierarchical clustering. This heat-map reflects the greater number of *C. albicans* genes that were up- and down-regulated in response to lactate at pH 5 than pH 7 (Figure 1B). Specific clusters were subjected to Gene Ontology (GO) analysis, revealing significant enrichment of GO terms relating to metabolism, and a substantial remodelling of the expression of genes involved in iron and zinc homeostasis (clusters 5 and 6). Significantly, genes such as *FET31*, *FTR2*, *ZRT101*, *PRA1*, *CTR1*, *FRE7,* and *FRE30* were induced by lactate at pH 5, but not at pH 7. On the other hand, *FTH1*, *FTR1*, *FET34*, and *CFL5* were repressed at pH 5, but not at pH 7.0. The *FRP1*, *FRE10*, and *FET99* genes were repressed by lactate at both pH 5 and 7. We did not detect changes in iron and zinc homeostasis genes at pH 7 that were not also observed at pH 5. These data suggested that, under the growth conditions analysed, exposure to lactate at pH 5 affects iron and zinc assimilation and homeostasis, but not at pH 7. Figure 1C shows the fold changes in expression of genes related to iron, copper, and zinc homeostasis (clusters 5 and 6) upon shifting to lactate-containing versus lactate-free media. An immediate response was detected after 30 minutes, followed by a general attenuation after one hour, which recovered at later time points. These data suggested that lactate influenced the homeostasis of these transition metals, particularly iron, we further confirmed and investigated this effect.

### Interplay between PKA signalling and iron homeostasis in the lactate-induced transcriptional response

To better understand the regulatory pathways controlling the observed transcriptional responses to lactate, we used the YEASTRACT+ database. YEASTRACT+ is a curated repository of regulatory associations between transcription factors and target genes in yeast species, with dedicated databases for 11 yeast species, including *C. albicans*. Specifically, with regards to *C. albicans*, YEASTRACT+ compiles over 50,000 regulatory associations between 129 characterized transcription factors and their target genes, as well as 93 defined transcription factor consensus sequences, based on curated published experimental results (22).

Using the *Rank by Transcription Factor* tool associated with YEASTRACT+, we identified transcription factors known to regulate most genes in the provided gene list (Table S5). Interestingly, this analysis identified Efg1 as the master regulator of the lactate-induced response, controlling 91 % of the 131 differentially expressed genes. The transcription factor Efg1, is regulated by the cyclic AMP/PKA pathway (23) and plays a key role in the control of hyphal growth, adhesion, biofilm formation and virulence (24–29). Interestingly, other PKA pathway-related transcription factors, namely Flo8 and Wor1, are predicted to control 56 % and 21 % of the lactate-induced response.

Other transcription factors that stood out in our analysis included a group of regulators involved in iron homeostasis, particularly Hap43 and Sfu1 (30). According to this YEASTRACT+ analysis, Hap43 and Sfu1 regulate 21 % and 8 % of the genes we observed in the lactate response, respectively (Figure 2A). This observation reinforced the outputs of the GO clustering (Figure 1B). Hap43 is a transcriptional regulator required for the response to low iron, whereas Sfu1 is a repressor of iron acquisition under iron-replete conditions. Additional transcription factors related to iron homeostasis include Aft2, Sef1, Hap2, Hap3, Hap5, and Crz1, which also control small subsets of the observed response (Figure 2A). Genes involved in iron acquisition are also positively regulated by the pH-responsive transcription factor Rim101 in neutral-alkaline environments (31) Significantly, our YEASTRACT+ analysis revealed that Rim101 controls 72 % of the lactate- induced response.

**Figure 2.**
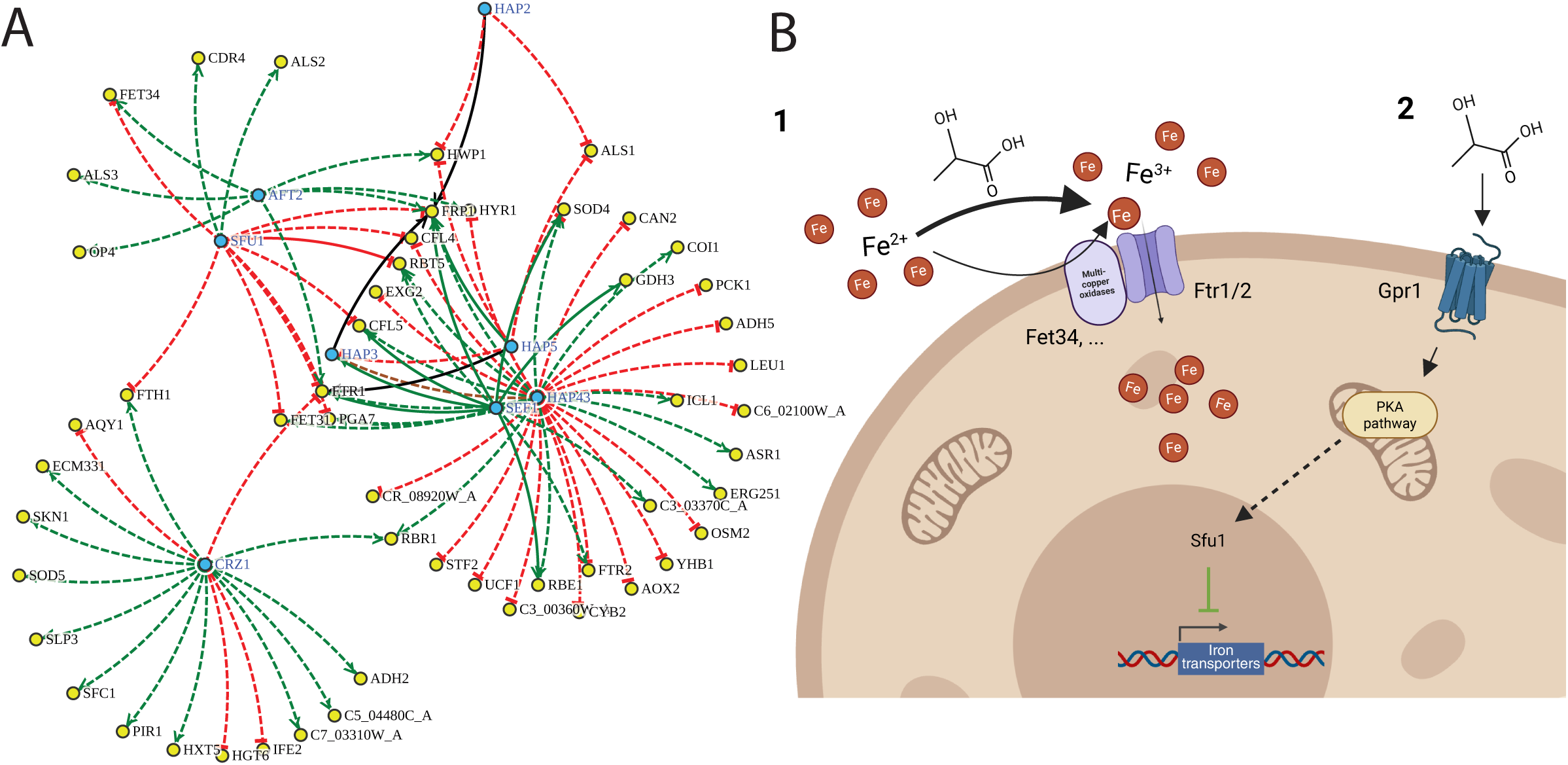
Iron-related transcription factor network predicted to be involved in the control of lactate-induced transcription remodelling. **(A)** This image was generated by the YEASTRACT+ database Rank by TF tool, upon selection of the subset of iron-related transcription factors, regulating the lactate-induced transcriptional response observed in this study. Full lines indicate regulatory associations known to be direct, based on transcription factor-DNA interaction experiments, while dashed lines indicate regulatory associations observed based on expression studies alone. Green and red lines indicate whether the transcription factors are known to act as activators or repressors of the corresponding target genes, respectively. **(B)** Illustrative scheme depicting the two major ways lactate might alter iron availability through either bypassing multicopper oxidases (for example Fet34) *via* lactate-dependent iron oxidation which is then transported by the high affinity iron permeases Ftr1/2 (1) or lactate-signalling through Gpr1 and the PKA pathway, ultimately altering the iron regulon (of which Sfu1, repressor of ferric uptake, is depicted), resulting in downregulation of iron transporters (2).

Taken together, our GO term and YEASTRACT+ analyses strengthened our working hypothesis that lactate might affect iron homeostasis in *C. albicans* and that PKA signalling might play an important role in regulating this lactate response.

We reasoned that iron oxidation by lactate might reduce the requirement for multicopper oxidases, resulting in a higher bioavailability through an enhanced iron reductive uptake pathway (option 1 in Figure 2B). Alternatively, a potential interplay between the PKA pathway and the iron regulon has been proposed (option 2 in Figure 2B), involving the G-protein-coupled receptor Gpr1, where the PKA pathway activates the kinase Ssn3, which in turn phosphorylates Hap34 a key transcriptional repressor involved in low-iron response, targeting it for degradation (32, 33). This post-translational modification alleviates the repression of Sfu1, a downstream regulator that controls iron acquisition machinery. This ultimately leads to the transcriptional downregulation of iron uptake systems, such as *FTR1/2* and *FET34*, which is consistent with our transcript profiling outputs following lactate exposure.

### Lactate promotes intracellular iron accumulation in an Ftr1-dependent manner

With a view to testing these working hypotheses (Figure 2B), we focused on *FTR1* and *FET34*, which encode a high-affinity iron transporter and a multicopper oxidase involved in iron oxidation and uptake, respectively (18, 34). qRT-PCR was performed to quantify the fold changes in these transcripts in the presence or absence of lactate. Fold changes were expressed relative to the zero time point without lactate (Figure 3A). These experiments confirmed the increases in *FTR1* and *FET34* expression between 0 and 30 min observed by RNA sequencing. *FTR1* and *FET34* expression levels declined after 30 min, irrespective of whether lactate was present. The data also confirmed that the increase in *FTR1* expression is significantly attenuated in response to lactate (*p* = 0.0016). Therefore, lactate induced transient changes in the expression of the key iron assimilation gene To evaluate whether these lactate-induced transcriptional changes affect iron uptake through the iron reductive pathway, we first compared intracellular iron levels in *C. albicans* wild-type and *ftr1Δ/Δ* strains (Figure 3B). In wild-type cells, we observed a significant increase in intracellular iron upon lactate supplementation relative to the control without lactate (*p* = 0.0094 and 0.0024 at 1 h and 4 h, respectively), suggesting that lactate promotes iron uptake. Cells lacking *FTR1* still displayed an increase in iron uptake at 1 h (*p* = 0.0224), but this increase was transient, and iron uptake decreased at 4 h, unlike the wild-type control. We conclude that Ftr1 is required for the sustained increase in iron uptake in response to lactate (Figure 3B).

**Figure 3.**
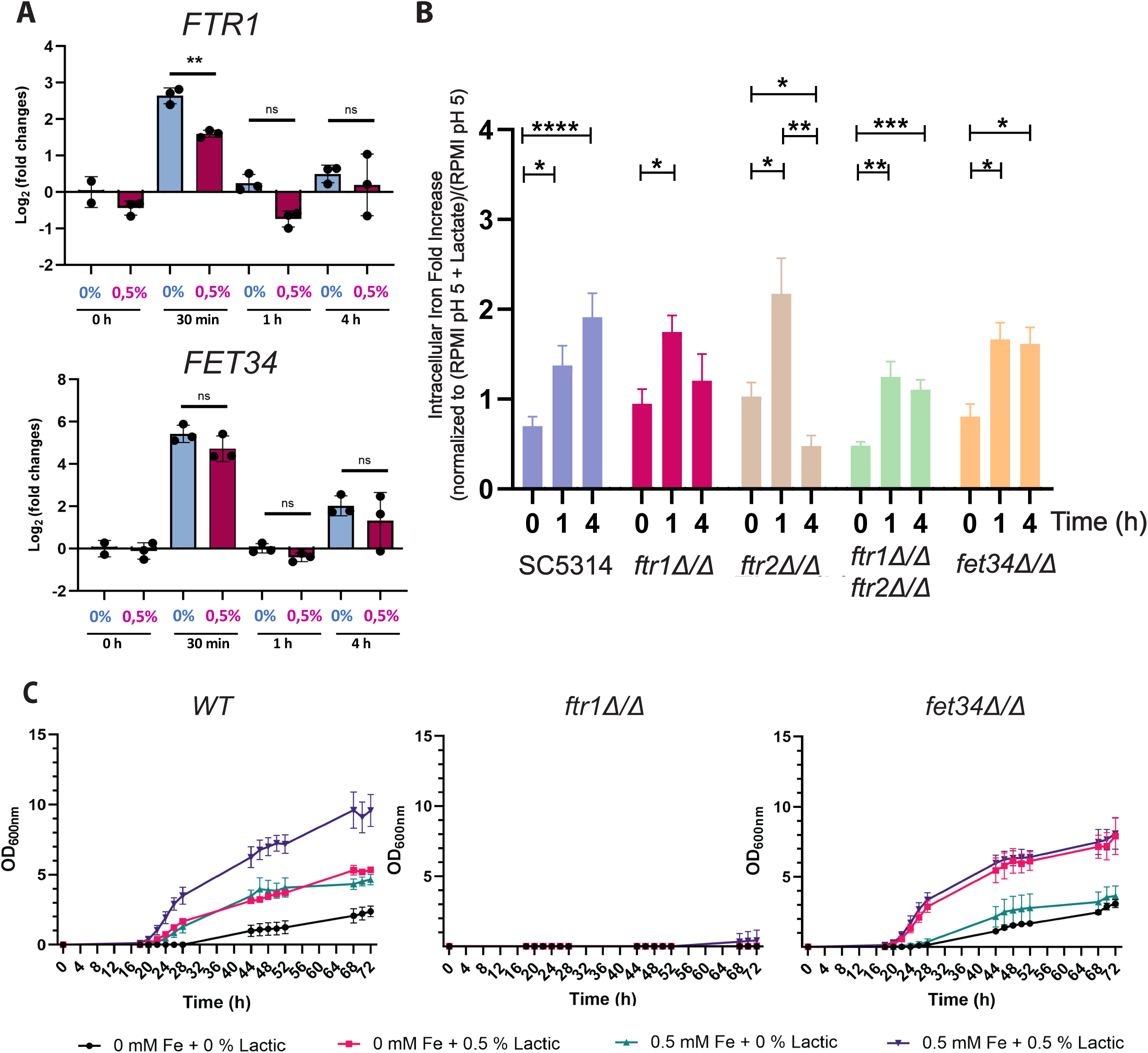
Lactate modulates the expression of iron transport genes and the intracellular iron content. **(A)** qPCR of *FTR1* and *FET34* expression levels in response to the presence of different concentrations of lactate at pH 5 over time. Graphs show fold variation of each gene compared to housekeeping genes *ACT1* and *TUB1*, scaled to the 0 % Lactate / 0-hours timepoint group. Statistical analysis was performed using a student’s t-test (n=3; *p < 0.05; **p < 0.01; “ns” indicates non-significance between conditions). Data are presented as mean ± SD from three independent biological experiments. **(B)** Intracellular iron fold variation in WT, *ftr1Δ, ftr1Δ, ftr1ftr2Δ* and *fet34Δ* strains. Values represent increases relative to RPMI pH 5 at the same timepoint. Statistical analysis was performed using a student’s t-test (*p < 0.05; **p < 0.01). Data are presented as mean ± SEM from four independent experiments obtained with four independent transformants. **(C)** Growth curves of the WT strain, *ftr1Δ* and *fet34Δ* mutant strains when grown in LIM supplemented with different lactate and iron concentrations. Cells were pre-grown in LIM, supplemented with 80 µM of BPS at pH 5 for twelve consecutive overnight cultures to starve them for iron. Lactic acid was tested at final concentrations of 0 % and 0.5 %, with iron (Fe^3+^) at final concentrations of 0 μM and 0.5 μM. Data are presented as mean ± SEM from three independent experiments.

We reasoned that *FTR2*, encoding another high-affinity iron permease that was upregulated in our study (Figure 1C), might partially compensate for the loss of *FTR1*. To test this, we examined a *C. albicans ftr2Δ/Δ* single mutant and a *ftr1Δ/Δ ftr2Δ/Δ* double mutant. These mutants behaved in a similar fashion to the *ftr1Δ/Δ* single mutant: iron uptake only increased transiently in response to lactate, in contrast to the sustained increase in iron uptake observed for wild-type control cells (Figure 3B). We then turned to the multicopper oxidase gene, *FET34.* Like the *ftr1Δ/Δ* and *ftr2Δ/Δ* mutants, *C. albicans fet34Δ/Δ* cells initially displayed an increase in intracellular iron after 1 h of lactate exposure, but unlike the wild-type control, no further increase was observed at 4 h. We conclude that Ftr1, Ftr2, and Fet34, and by implication the iron reductive uptake pathway, promote sustained iron assimilation in response to lactate (option 1 in Figure 2B). However, other factors appear to be required for the initial increase in iron assimilation in response to lactate.

We then evaluated the impact of lactate-enhanced iron uptake on the ability of *C. albicans* to grow under iron limitation, and whether lactate suppresses the need for the conventional reductive iron uptake pathway under these conditions. To achieve this, we grew wild-type, *ftr1Δ/Δ,* and *fet34Δ/Δ* cells under iron limiting conditions for 12 days and subsequently re-administered either lactate (0.5 %), iron (0.5 µM), or both in combination. Wild-type cells grew slowly in the absence of lactate and iron (Figure 3C; asymptotic maxima (A) and growth rates (µ) are presented in Supplementary Table 3; Statistics are presented in Supplementary Table 4). As expected, the addition of iron to the iron-limited medium enhanced growth. Interestingly, lactate significantly enhanced the growth of wild-type cells under iron limiting conditions, and growth was further enhanced by lactate even when iron was not limiting (Figure 3C, Supplementary Table 3). The *fet34Δ/Δ* mutant grew under iron limitation in the absence of lactate, possibly because other multicopper oxidases in *C. albicans* (Fet3, Fet31, Fet33, Fet99) compensated for the loss of Fet34. Interestingly, the *fet34Δ/Δ* cells grew well with lactate even in the absence of supplementary iron (Figure 3C, Supplementary Table 3). Meanwhile, as expected (34), cells lacking the Ftr1 high-affinity iron transporter were unable to grow under iron limiting conditions. The addition of lactate was unable to suppress this growth defect (Figure 3C, Supplementary Table 3). We conclude that lactate enhances the growth of *C. albicans* under iron limiting conditions, and that the sustained iron uptake mediated by Ftr1 is important for this effect (Figure 2B, option 1).

We then tested whether lactate-induced iron assimilation is dependent upon PKA signalling (Figure 2B, option 2). To achieve this, we exploited two inhibitors of PKA signalling: H-89, a specific inhibitor of PKA in *C. albicans* (35); and farnesol, which directly inhibits adenylyl cyclase on the PKA pathway (36). Neither inhibitor blocked the lactate-induced iron uptake observed for untreated wild type cells (Figure 4). We conclude that, despite the apparent involvement of Efg1 in the transcriptional response to lactate (above), lactate-induced iron assimilation does not depend upon PKA signalling, in the conditions tested.

**Figure 4.**
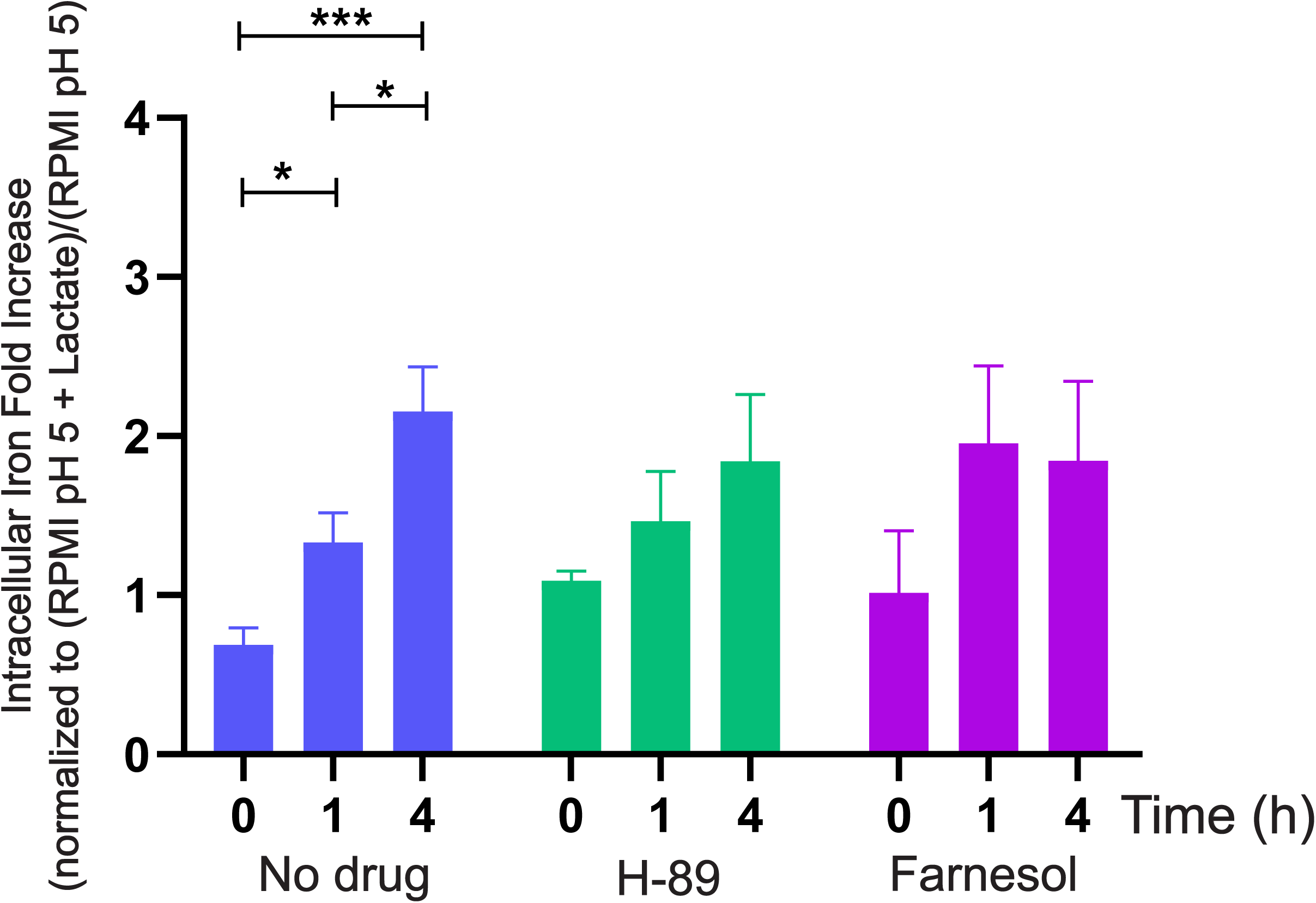
Inhibiting PKA signalling by H-89 or farnesol did not block the enhanced iron assimilation induced by lactate. Intracellular iron levels measured in wild-type *C. albicans* (SC5314) and cells treated with either 50 µM H-89 or 150 µM farnesol over 0, 1, and 4 hours. Values are expressed as fold increase in intracellular iron, normalized to the expression ratio of RPMI pH 5 + lactate over RPMI pH 5. Data represent the mean ± SEM of biological replicates (n=3). Statistical analysis was performed using student’s t-test (*p < 0.05; *p < 0.01; ***p < 0.001)

## Conclusion

Our genome-wide profiling of *C. albicans* cells growing on RPMI revealed that exposure to physiologically relevant concentrations of lactate influences the expression of genes involved in transition metal homeostasis. This effect, which included elements of the reductive iron uptake pathway, occurred at pH 5, but not at pH 7. At first glance, our observations might appear to contradict those of Cottier and coworkers (20), who reported that, in response to weak acids, *C. albicans* upregulates (rather than downregulates) genes encoding iron transporters and assimilates lower (rather than higher) levels of intracellular iron. However, the authors examined the effects of a relatively high concentration of lactate (0.62 M) on *C. albicans* cells grown in Man, Rogosa, Sharpe (MRS) medium, which they selected because this rich medium supports the growth of *Lactobacillus* (20).

At such high concentrations, lactate is thought to reduce iron bioavailability by counteracting the iron-liberating mechanisms of ferric reductases (17) . In contrast, our experiments employed ∼10-fold lower concentration (0.5 % = 0.055 M), which are within the physiological range. Under these conditions, lactate is more likely to promote Fe^2+^ oxidation, thereby complementing the activity of multicopper oxidases, without unduly interfering with the ferric reductases.

Indeed, we confirmed that *C. albicans* cells accumulate intracellular iron in response to this physiologically relevant lactate concentration and consistent with previous findings (37), our transcriptomic data showed that this increased iron availability led to downregulation of genes in the iron reductive uptake pathway.

Efg1 was also identified as a regulator of the observed transcriptomic response. This was consistent with the involvement of PKA signalling in lactate-induced changes in *C. albicans* physiology (7, 9, 38). However, pharmacological inhibition of PKA signalling by H-89 or farnesol did not block the enhanced iron assimilation induced by lactate. Instead, this sustained iron accumulation was dependent on the iron reductive uptake pathway (*FTR1*). Significantly, this sustained iron accumulation underlies the ability of lactate to stimulate the growth of *C. albicans* under iron limiting conditions. Given the importance of nutritional immunity in limiting microbial infection (39–41), this interaction between lactate and iron assimilation is likely to enhance fungal colonisation of host niches.

## Materials and Methods

### Yeast Strains and Growth Conditions

All *C. albicans* strains used in this study were derived from the clinical isolate SC5314 and are listed in Table S1. Strains were routinely cultured in yeast extract-peptone-dextrose (YPD) medium, at 30 °C. For RNA extraction, cells were grown in filter-sterilized RPMI 1640 medium (Sigma, R6504) buffered with 0.165 M morpholinepropanesulfonic acid (MOPS, Sigma) at the specified pH, unless otherwise stated. Growth experiments were performed on cells grown in filter-sterilized yeast nitrogen base with amino acids, containing 0.2 % (w/v) of glucose, without supplemented iron, and buffered with 0.165 M morpholinepropanesulfonic acid (MOPS).

### RNA extraction, sequencing, and analysis

*C. albicans* SC5314 cells were grown overnight in YPD at 30 °C. Cells were then harvested, washed twice in 1X phosphate-buffered saline (PBS), and transferred to 0.2 % (w/v) glucose RPMI 1640 at pH 5 or 7 (30°C) at an initial OD600 of 0.3. When cultures reached an OD600 of 1.0, sodium lactate was added to a final concentration of 0 or 0.5 % (v/v) (t = 0). The *C. albicans* cells were harvested by centrifugation at 0, 0.5, 1, 2, and 4 h after lactate addition, and stored at -80 °C for RNA extraction.

To extract RNA, cells were resuspended in 0.5 mL of extraction buffer (1.0 mM EDTA, 0.1 M LiCl, 0.1 M Tris-HCl pH 7.5) and 0.5 mL of phenol-chloroform-isoamyl alcohol (PCI) containing 1 % sodium dodecyl sulphate (SDS). The samples were then homogenized with zirconium beads using a FastPrep-24™, centrifuged at 14,000 rpm for 10 minutes at 4 °C, and the aqueous upper layers transferred to clean Eppendorf tubes. The PCI extraction, using PCI without SDS, was repeated until no interphase was visible. Subsequently, 1/10 volume of 40 % potassium acetate, pH 5.5, and 2 volumes of 100 % ethanol were added, and the samples were mixed and stored at –20 °C. Samples were centrifuged at 14,000 rpm for 10 min at 4 °C, the pellets were washed with 0.5 mL of 70 % ethanol, and dissolved in 50 µL of RNAse-free water. RNA concentrations were measured using a NanoDrop® ND-1000, with an RNA quality cut-off of OD260nm /OD280nm > 2.

Library preparation and RNA sequencing were performed by the University of Exeter Sequencing Facility. Libraries were prepared with the NEBNext® Ultra II Directional RNA Library Prep Kit (NEB #E7760, New England Biolabs) after poly(A) enrichment. Paired-end sequencing was performed in an Illumina NovaSeq SP (150 nt reads). Sequence reads were mapped against the *C. albicans* SC5314 genome (assembly ASM18296v3 using STAR version 2.7.9a). Expression levels were quantified using RSEM (version 1.3.1) and normalised using the variance stability transform in DES2 (version 1.38.3)(42). The Benjamini and Hochberg’s approach was used to adjust the resulting *p*- values (43), and genes with an adjusted *p*-value <=0.05 and a log (2) fold change of +/- 1 were assigned as differentially expressed. Gene Ontology analyses for functional profiling were done with the g:Profiler server (https://biit.cs.ut.ee/ gprofiler/gost) using *C. albicans* SC5314 data and default settings.

The YEASTRACT database (22) was employed to identify potential master regulators underlying the *C. albicans* response to lactic acid. Using the *Rank by TF* tool, those transcription factors that are known to control the genes whose transcription was found to be affected by lactate were identified and ranked according to the percentage of targets in the provided gene list (Table S5). The ranking also took into consideration the enrichment of each transcription factor regulon in the provided gene list, and computed that enrichment in terms of *p*-value, calculated using a hypergeometric test.

### Quantitative Real-Time Reverse Transcription PCR (qRT-PCR)

cDNA was prepared using the iScript™ cDNA Synthesis Kit (Bio-Rad Catalog 170-8891) in a 20 µL reaction containing 2 µg of total RNA, according to the manufacturer’s instructions. Real-time quantitative PCR was performed on these cDNA samples using the GoTaq® qPCR kit on a StepOnePlus™ Real-Time PCR System. Primers and thermocycling conditions are described in Supplementary Table 2. Reaction mixtures were set up in a total volume of 20 μL using 10 μL of GoTaq® qPCR Master Mix 2x, 0.2 μL of CXR Reference dye (30 μM), 10 μM of each primer, 5 μL of cDNA (0.4 ng/μL), and nuclease-free water. A negative control without template was used for each target transcript in each run. Expression levels were normalised against the *ACT1* and *TUB1* internal controls.

### Determination of iron concentration

Intracellular iron levels were quantified using the BPS-based colorimetric method (44). Briefly, *C. albicans* cells were grown as described above for RNA extraction. *C. albicans* cultures were harvested at an OD600 of 1.0. Cell pellets were washed with ddH2O, resuspended in 500 μL of 3% nitric acid (Fluka Analytical), and boiled for 2 h at 98 °C. Cell debris was then removed by centrifugation. Four hundred µL of the resulting supernatants were mixed with 160 µL of 38 mg/ml sodium ascorbate (Sigma-Merck), 320 µL of 1.7 mg/ml BPS (Sigma), and 126 µL of 4 M ammonium acetate (Merck). These reaction mixtures were incubated at room temperature for 15 minutes, and the OD535 was measured with a spectrophotometer (Synergy H1, BioTek) against blanks containing all reagents except cells. Nonspecific absorbance was removed by subtracting the OD680 from the OD535. Values were plotted against a calibration curve (0, 1, 5 & 10 µM of Fe(II)SO4.7H2O) to determine the iron concentration.

### Growth Analysis

The growth of *C. albicans* SC5314 WT, *ftr1Δ*, and *fet34Δ* strains was examined under iron limiting conditions. Cells were pre-grown in 1.5 ml of Limiting Iron Medium (LIM) at pH 5 (6.9 g/L YNB without Iron (CYN1102, Formedium), 0.79 g/L CSM mix (MP Biomedicals); 0.2 % (w/v) glucose, supplemented with 80 µM of BPS) for twelve consecutive overnight cultures. A 1:100 dilution was made after the first day, and for the remaining 11 days, the entire overnight cell culture was harvested, and the cells were transferred to fresh medium. For each strain, the 12-day starved pre-culture was collected, harvested, and washed once with 1 mL of LIM. The cells were then transferred to 50 mL of growth medium in 300 mL flasks, with an initial OD600 nm of 0.00001. Four different media were tested: 0 versus 0.5 µM of FeCl3, and 0 versus 0.5 % (v/v) lactate, supplemented with 10 µM of BPS, at pH 5.0. To reduce noise and improve growth fitting stability, replicates for each condition were averaged at each time point. The resulting averaged growth curves were fitted to the Gompertz growth model, which estimates the asymptotic maximum (A), the growth rate (µ), and the lag time with their respective standard deviation. These SDs reflect the sensitivity of the model to the observed data and are used for subsequent statistical inference.

To assess statistical differences in growth dynamics, Z-tests were conducted by comparing each condition to a baseline condition (0_0). To account for multiple hypothesis testing, *p*-values were corrected using the Benjamini-Hochberg procedure to control the false discovery rate (FDR). This correction was applied separately for each growth parameter across all pairwise and baseline comparisons. Adjusted *p*-values below 0.05 were considered statistically significant.

### Statistics and reproducibility

Statistical analyses were performed in Graphpad Prism v10 using Two-way ANOVA and Student’s t–test or the Python packages SciPy with Statsmodels and a Z-test. Differences were considered significant at P≤0.05.

### Data availability

RNA-Seq data was deposited at *Repositório de Dados da Universidade do Minho* (dataRepositóriUM) under study https://doi.org/10.34622/datarepositorium/LWNKLQ

## Acknowledgements

We are grateful to Karen Moore, Jemima Onime and Paul O’Neill in the Sequencing Facility at the University of Exeter for their help with the generation and analysis of the RNA sequencing dataset, which was performed using equipment funded by Wellcome (218247/Z/19/Z).

This work is supported by the MetaFungal project PTDC/BIA- MIC/5246/2020 (https://doi.org/10.54499/PTDC/BIA-MIC/5246/2020), funded by the Portuguese Foundation for Science and Technology. Work at the University of Minho (CBMA) was supported by the ‘Contrato-Programa’ UIDB/04050/2020 (https://doi.org/10.54499/UIDB/04050/2020) and the Contrato-Programa” LA/P/0069/2020 (https://doi.org/10.54499/LA/P/0069/2020), funded by national funds through the FCT I.P

Work at KU Leuven (Laboratory of Molecular Cell Biology) was supported by a grant from the Fund of Scientific Research Flanders (FWO # G0C0622N) and by a grant from the KU Leuven Research Fund (# C14/22/075) AG-G acknowledges FCT for the 2021.08564.BD PhD grant (https://doi.org/10.54499/2021.08564.BD).

WVG was supported by the KU Leuven Research Council (grant # C14/22/075).

AJPB was supported by a programme grant from the UK Medical Research Council [MR/M026663/2], a Wellcome Investigator Award (224323/Z/21/Z), the Medical Research Council Centre for Medical Mycology at the University of Exeter (MR/N006364/2 and MR/V033417/1), and the NIHR Exeter Biomedical Research Centre. The views expressed are those of the authors and not necessarily those of the NIHR or the Department of Health and Social Care. For the purpose of open access, the author has applied a CC BY public copyright licence to any Author Accepted Manuscript version arising from this submission.

JA was supported by grants PID2020-113319RB-I00 and PID2023-150535OB-I00 (Ministerio de Ciencia, Innovación y Universidades, and Agencia Estatal de Investigación, Spain).

The funders had no role in study design, data collection and analysis, decision to publish, or preparation of the manuscript.

---

Figure S. 1- Transcriptional impact of lactate and pH in C. albicans Venn diagram showing the number of genes differentially induced at least at one time point in cells cultured in media with or without lactic acid at pH 5 and 7.

**Figure S1.**
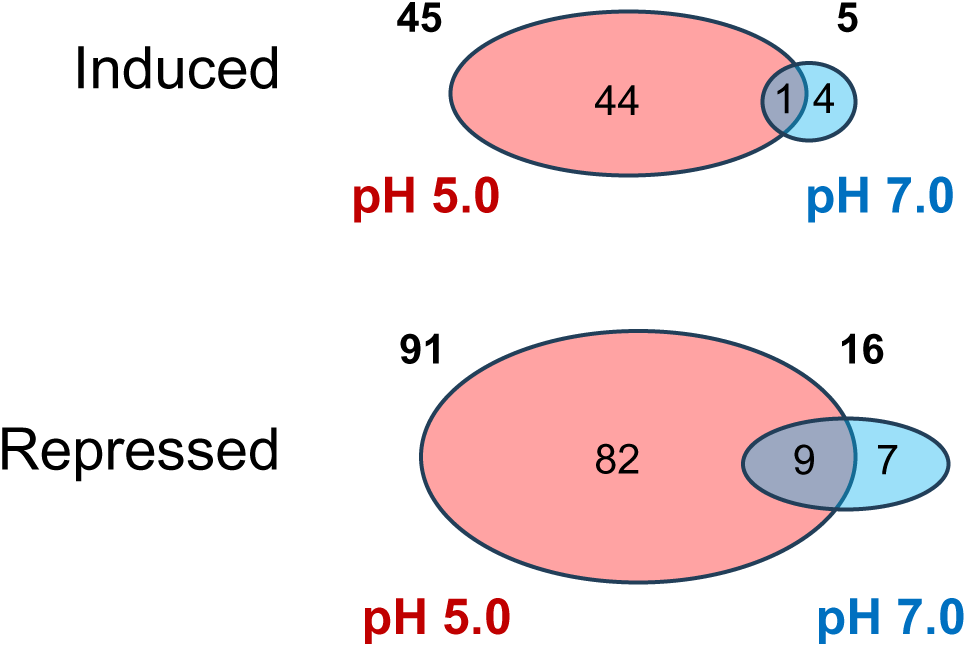
Venn diagram showing the number of differentially induced genes at least one time point in cells cultured in the presence or absence of lactate in the medium at pH 5.0 and 7.0.

